# Co-evolved maternal effects selectively eliminate offspring depending on resource availability

**DOI:** 10.1101/2020.07.30.228528

**Authors:** Bin-Yan Hsu, Martina S. Müller, Christoph L. Gahr, Cor Dijkstra, Ton G. G. G. Groothuis

**Author notes:** deceased (Cor Dijkstra).

## Abstract

Many plants and animals adaptively downsize the number of already-produced propagules if resources become insufficient to raise all of them. In birds, mothers often induce hatching asynchrony by incubating first eggs before last eggs are laid, creating an age/size hierarchy within broods which selectively eliminates the smallest chicks in poor food conditions. However, mothers also deposit more testosterone into late-laid eggs, which boosts competitive abilities of younger chicks, counteracts the competitive hierarchy, and ostensibly creates a paradox. Since testosterone also carries costs, we hypothesized that benefits of maternally deposited testosterone outweigh its costs in good food conditions, but that testosterone has a net detrimental effect in poor food conditions. We found experimental evidence that elevated maternal testosterone in the egg caused higher chick mortality in poor food conditions but better chick growth in good food conditions. These context-dependent effects resolve the paradox, suggesting co-evolution of two maternal effects, and explain inconsistent results of egg hormone manipulations in the literature.

## Introduction

Throughout the plant and animal kingdoms, mothers routinely overproduce propagules, and competition between them culls superfluous offspring until family size matches available resources^1,2^, thereby neatly (albeit harshly) resolving the evolutionary trade-off between offspring quantity and quality. The severity of resource limitation determines the intensity of propagule competition; when food is scarce, siblings die, e.g. by siblicide in spotted hyena broods^3,4^ and blue-footed booby chicks^5^, intra-fruit seed abortion in plants^6^, maternal crushing and/or starvation in piglets^7,8^, or sibling cannibalism in ladybird beetles^9^. Mothers can improve the efficiency with which sibling rivalry eliminates offspring by creating age/size hierarchies among their young that make some siblings less able to compete than others, allowing efficient culling of their numbers when needed. For example, in birds, mothers induce hatching asynchrony by initiating incubation before all eggs have been laid. This causes chicks from later-laid eggs to have a later start on development, to hatch later, and therefore to be smaller and weaker when competing with their older siblings for parental food provisioning.

Mothers also employ other tools for creating competitive asymmetries, including furnishing their embryos with maternal hormones^10^. Such hormone-mediated maternal effects have substantial and long-lasting impacts on the development of morphology, brain and behaviour^10,11^ and are important in plants^12^, insects^13^, reptiles^14^, fish^15^ and other vertebrates^10,16^. For example, in spotted hyenas, cubs from high-ranked mothers are exposed prenatally to higher levels of maternal androgens via the placenta, which makes them more aggressive after birth^17^. In birds, maternal androgens, like testosterone (a potent sex steroid hormone) are deposited into eggs in substantial quantities that vary systematically with environmental conditions and also according to the position of the egg in the laying sequence in a given breeding attempt (i.e. clutch)^18,19^. Because in most bird species, later-laid eggs contain higher concentrations of maternal androgens, the hatching asynchrony adjustment hypothesis (HAAH)^18,19^ proposes that this increased deposition of maternal androgens in late-laid eggs functions as a compensation for the competitive disadvantage of the chicks from these eggs. Indeed, experimental manipulation of egg androgens, especially testosterone, has revealed that these hormones stimulate the competitive ability of the chick and this has become a well-cited example of hormone-mediated adaptive maternal effects^19-23^.

The avian system of hatching asynchrony and maternal egg yolk hormones thus provides an excellent study model for the interactions of two maternal effects. It also reveals a significant paradox that has not yet been addressed adequately: if hatching asynchrony itself is an adaptive mechanism to achieve efficient brood reduction by sibling competition depending on the amount of food available^1,24^, why would mothers counteract this by increased androgen deposition in the later-laid eggs^25,26^?

As the research effort in this field grows rapidly, evidence that maternal androgens in egg yolks do not always counteract the effects of hatching asynchrony has accumulated. An increasing number of studies also found inconsistent effects from the experimental elevation of egg androgen concentrations, with sometimes opposite effects in the same species^23^. Rather than raising doubt about the adaptive explanations on within-clutch difference of maternal androgens, these inconsistent effects open up the possibility for an intriguing potential resolution to the HAAH paradox. Yolk androgen has been demonstrated to have both beneficial and detrimental effects. Thus, yolk androgens may have different effects depending on the food availability, and the androgens may act in concert with the effects of hatching asynchrony, which are also dependent on food availability. Egg androgens and hatching asynchrony together may then facilitate chick survival in good food conditions and promote brood reduction in poor food conditions.

Yolk androgen provides important benefits by increasing competitive ability in chicks^23^, but it can also impose significant costs, such as increased resting metabolic rate in the short- and long-term^27-29^, suppressed immune function^30-34^ and oxidative damage^35-39^. We expect higher maternal androgen to undermine survival in a year when resources are insufficient to offset its costs, but in a year when food is abundant, higher maternal androgen exposure may help chicks obtain enough food to offset the higher energy expenditure and the challenge to their immune system, and help them survive.

We therefore hypothesize that androgens in eggs promote adaptive resource-dependent brood reduction^26^. If so, two pathways have co-evolved that allow mothers to facilitate resource-dependent brood reduction: in poor-food years, one pathway culls the brood to preserve offspring quality, and in good-food years, the other pathway promotes survival to maximize offspring quantity. Combined, these two pathways maximize the reproductive value of the brood in all food conditions. The key to testing these resource-dependent effects of egg yolk androgens would be to combine a manipulation of yolk androgens at the time of egg-laying, mimicking elevated maternal androgen deposition, and a manipulation of food availability during the chick rearing phase, in a full factorial design. To our knowledge, no such experiment has been conducted to test our hypothesis.

We performed this experiment in rock pigeons (*Columba livia*), an excellent species for our study for two reasons: (1) its modal clutch size is two^40^ with the second egg always containing much higher amounts of testosterone in the yolk than the first one^41,42^, and (2) it displays substantial hatching asynchrony and the second chick is always smaller than the first one. In this study, we created experimental clutches of first-laid eggs only, which contain low levels of maternal testosterone. One egg was injected with testosterone solution to increase testosterone levels to those of second-laid eggs, and the other was injected with vehicle as control. Both eggs were given to foster parents and half of the resulting broods were reared in experimentally-imposed poor food conditions while the other half were raised in good food conditions. Over the nestling period, we monitored various aspects of growth and development as well as immune function and survival of the chicks. We expected to see beneficial effects of elevated yolk testosterone on chick growth and immune function under the good food condition, and detrimental effects under the poor food condition, specifically, increased early mortality of chicks.

## Materials and Methods

We used pigeons from our rock pigeon colony housed in a large outdoor aviary. Before the experiment started (early April, 2012), breeding pairs of adult pigeons were re-housed in smaller identical aviaries, with two pairs in each aviary (see Supplementary Material for housing details). All experimental and animal care procedures were under the approval of the animal welfare committee of University of Groningen (DEC No. 5635D) and complied with Dutch law.

### Egg collection, incubation, hatching time and the creation of experimental broods

In May (the annual peak of egg laying), we made the nest boxes available to induce breeding. We checked nest boxes every morning and marked and collected all freshly-laid eggs, and replaced with a dummy egg so parents would start incubating. To increase the sensitivity of our experimental design, we aimed at creating experimental broods consisting of chicks of opposite treatments (testosterone or control injection) that were matched as much as possible in body mass, sex and hatching time. We only used first-laid eggs (1^st^ eggs) in the experiment. Collected eggs were stored in a climate cell with relatively constant temperature (12-16°C) and humidity (40-50%) for no more than three days, until a large enough batch of eggs had been collected for creating experimental pairs. We paired eggs by matching laying date and mass as closely as possible, and then injected one egg in each pair with testosterone dissolved in sesame oil that increased levels in these 1^st^ eggs up to the level of the 2^nd^ eggs (testosterone eggs, see *Egg injections* in Supplementary Material) and injected the other with sesame oil only (control eggs). After injection, these eggs were returned immediately to an unrelated foster nest for incubation. The mass of testosterone-eggs and control-eggs did not differ significantly (mean±SD: testosterone-eggs, 17.00±1.27 g, n=95; control-eggs, 17.07±1.38 g, n=95; t=0.383, p=0.7021), nor laying date (Mann-Whitney U test, p=0.9779).

On day 16 of incubation (about two days before hatching), all eggs were placed into an incubator until hatching and replaced in the nest by dummy eggs. The incubator was maintained at a constant 37.5 °C with humidity > 75%. We checked the incubator every four hours between 9 am and 9 pm. We estimated hatching of hatchlings found at 9 am by the dryness of the down feathers as either at 3 am (with relatively dry down) or at 7 am (with wetter down). We measured body mass, head-bill length, tarsus length, and wing length of each hatchling, and took a small blood sample (< 75μl) from the medial metatarsal vein for sexing. We sexed the hatchlings with a molecular sexing procedure following the protocol described in Goerlich et al. (2009, 2010)^41,43^. We then matched hatchlings with opposite egg injection treatments in pairs by their hatching time and body mass, and returned paired hatchlings to unrelated foster nests. We successfully created 19 same-sex testosterone-control pairs of chicks with no significant difference in hatching time (n=29, t=-1.79, p=0.084) or body mass (t=-1.229, p=0.229). Ten pairs were fostered by parents under good food conditions and 9 pairs by parents under poor food conditions.

### Food treatment

We initiated the food treatment for parents on day 16 of incubation. Prior to the food treatment, a commercial seed mixture (KASPER™ 6721 + 6712, Table S1), water, and a mixture of small stones and pigeon grit was available *ad libitum* to all pigeons. From the start of the food treatment onwards, pigeons in the good food condition group were also additionally provided with *ad libitum* pigeon pellets (KASPAR™ P40, Table S1) and supplemented with vitamin powder (Supralith™). The food availability of the pigeons in poor food conditions was limited to 33 g of the seed mixture with broken corn kernels (lower protein and fat content, Table S1) per pair per day, the amount of average daily food consumption by a pair of adult pigeons according to our previous measurements. To accommodate increased energy demands as chicks grew, we provided the food-restricted group with additional food when chicks were older than one week, based on Jacquin et al. (2012)^44^. For every 7-day-old or older chick, we provided 8 g additional food; for every 14-day-old or older chick, 16 g of additional food was provided.

After the experiment, the body mass loss in adult pigeons in poor food conditions was significantly more than that of adults in good food conditions [good food condition, mean (% of body mass before egg-laying)±SE = −6.31±0.75, n=67; poor food condition, mean±SE = −13.77±1.16, n=45, t=5.395, p<0.0001], indicating that our experimental food restriction significantly reduced energy intake to the adults.

### Chick monitoring

Nests were checked every day to monitor chick survival. Four biometric variables: body mass, head-bill length, tarsus length, and wing length were measured every two days in the first two weeks after hatching, and every three days from two weeks to 26 days post-hatching. All body measurements were taken by the same experimenter (the first author). We use SRBC test and hemagglutination assay to measure the humoral immune response of the experimental chicks shortly after fledging (37-45 days old). Pre-treatment blood samples were taken right before SRBC inoculation. Post-treatment samples were taken six days later. See supplementary material for details.

### Statistical analysis

All statistical analyses were performed with the software R 3.5.2^45^. We used a binomial linear model to test the effects of treatment on hatching success, with hormone injection and egg mass as predictors. We performed a Mann-Whitney U test to test for effects on hatching time, because it was not normally-distributed (Fig. S1). We analysed hatchling body mass using a general linear model (GLM), including hormone injection, egg mass, and sex as predictors and also tested for sex-specific effects of prenatal testosterone on hatchling body mass by including the interaction between hormone injection and sex.

We only tested the effects of treatment on chick body measurements until day 8, because the sample size for testosterone chicks in the poor food condition after day 8 decreased drastically due to the high mortality in this group (see *Results*), yielding low statistical power and potential computational problems if analysing the whole growth curve. In addition, for chicks reared in good food conditions we also tested the effect of testosterone on each body measurement at day 26, just before fledging. We used a general linear mixed model (R package *lme4*)^46^, and Kenward-Roger approximation method to compute p-values (R package *pbkrtest*)^47^. We included brood identity as a random factor, and hormone treatment, food treatment, and sex as fixed factors. We also included egg mass as a covariate, as it is known to strongly influence nestling body mass^48^. The interactions between hormone and food treatment (for day 8 only), and between hormone treatment and sex (for both day 8 and day 26) were also tested. Significant interactions were then further analysed with the package *phia*^49^ for pair-wise comparisons (Holm-adjusted p-values are presented). The results of the four biometric variables showed clear consistency with each other, with effects on body mass being the most pronounced. Therefore, only the results of growth in terms of body mass are reported below and the other three (head-bill, tarsus and wing length) are reported in the supplementary materials.

For our analyses of immune response, we subtracted for each chick the pre-treatment score from the final score of the SRBC-hemagglutination test. We applied a GLM to the resulting values, with hormone injection, food treatment, and sex included as independent variables. The model residuals did not violate assumptions of normality (Shapiro-Wilk test, p=0.492). The interactions of hormone injection with sex and hormone injection with food were also tested.

Chick survival was assessed between hatching to day 26, yielding right-censored data. Kaplan-Meier survival curves for pigeons hatched from the testosterone-injected and control eggs and reared under good and poor food conditions were built and tested using the package *survival*^50^ with Gehan-Wilcoxon test and rho = 1.

## Results

### Hatching success, hatching time, and hatchling body mass

We injected a total of 190 eggs (95 testosterone-injected eggs and 95 control eggs) of which 93 eggs hatched successfully (48.95%). Of those 93 eggs, 44 were control eggs and 49 were testosterone-injected eggs and this difference was not significant (n=190, p=0.407). Heavier eggs were more likely to hatch successfully (estimate±SE = 0.338±0.117, z=2.895, p=0.004). There was no difference in time of hatching between testosterone-injected and control eggs (Mann-Whitney U test, W=1080, p=0.991).

Of the 93 hatchlings, 90 had no visible developmental abnormalities and were included in the analysis of hatching body mass. Testosterone treatment did not affect hatchling body mass (marginal means±SE: hatchlings from testosterone-injected eggs, 11.58±0.11; control hatchlings, 11.78±0.11, t=-1.291, p=0.200) and hatchling body mass was also not affected by sex (marginal means±SE: males, 11.66±0.11; females, 11.70±0.11, t=-0.281, p=0.780) or the interaction between sex and hormone treatment (p = 0.971). Only egg mass significantly predicted hatchling body mass (estimate±SE = 0.612±0.060, t=10.152, p<0.001).

### Chick survival

Among the 90 hatchlings, we successfully created 19 same-sex testosterone-control pairs of chicks with 10 pairs reared under good food conditions and 9 pairs under poor food conditions. Their survival rates differed significantly among the four combinations of food treatment and testosterone injection (p<0.001, Fig. 1). In the good food conditions, chick survival was 100% (n=20), regardless of hormone treatment, while in the poor food conditions, chicks from testosterone-injected eggs (hereafter “testosterone chicks”) had lower survival than chicks from control eggs (hereafter “control chicks”) did (n=18, χ^2^=3.9, p=0.049, Fig. 1). Overall, chicks reared under the poor food conditions had significantly lower survival than those under the good food conditions (n=38, χ^2^=12.8, p<0.001).

**Figure 1.**
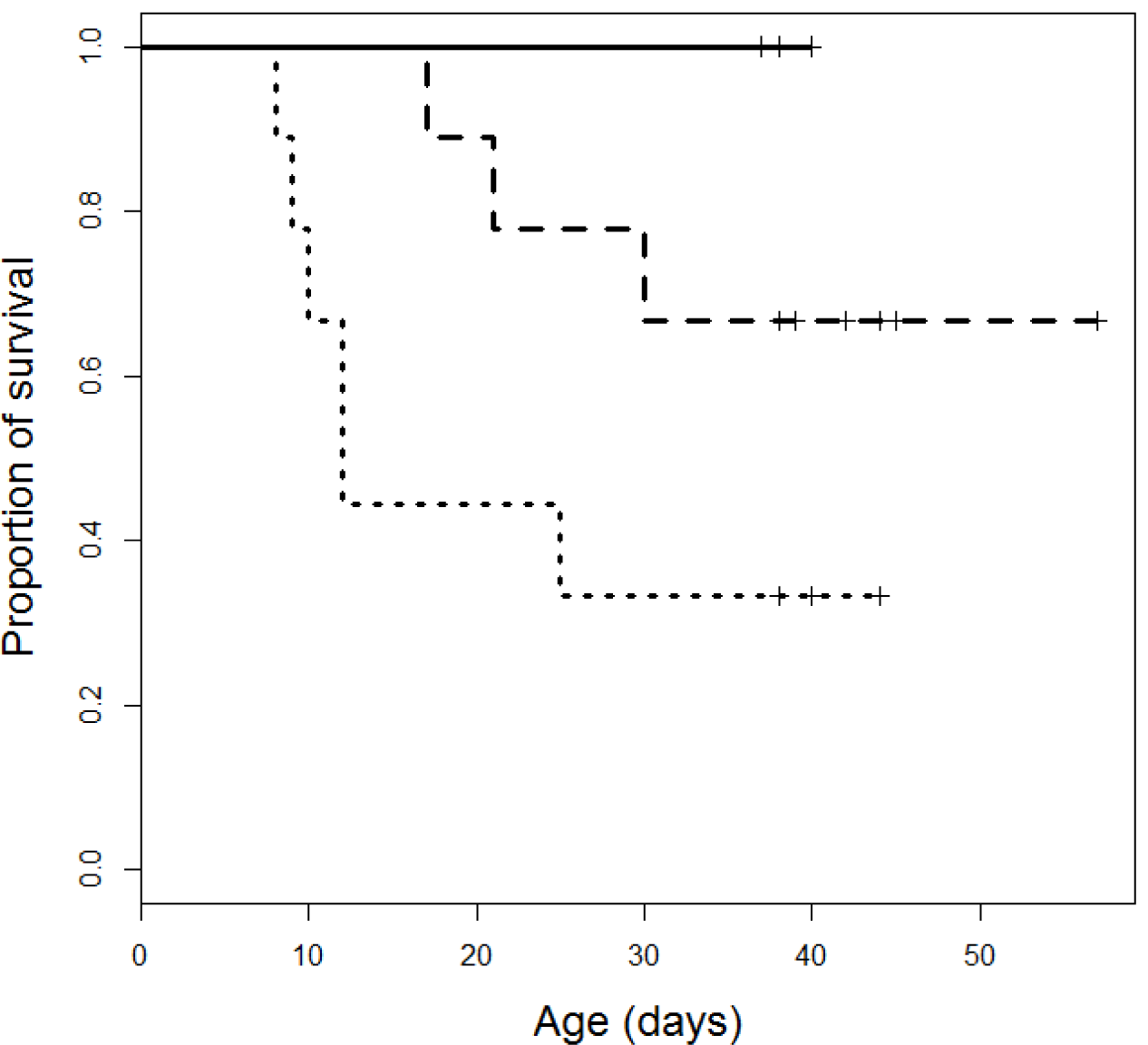
Survival curves of testosterone- and control-chicks. In good food conditions (solid line), chicks from both egg injection treatment had 100% survival so curves for testosterone- and control-chicks overlap completely. In poor food conditions, Chicks from testosterone-injected eggs (dotted line) had lower survival than control chicks (dashed line).

### Body mass at day 8 and day 26 after hatching

For day-8 body mass, the mixed model showed that the interaction effect of testosterone injection and food treatment was significant (F_1,16.66_=6.758, p=0.019, Fig. 2). Post-hoc interaction analysis indicated that testosterone chicks were heavier (marginal mean±SE = 120.49±4.42 g) than control chicks (marginal mean±SE = 103.69±4.42 g) in the good food conditions (χ^2^=10.577, Holm-adjusted p=0.002), but not in the poor food conditions (marginal means±SE = 55.74±4.69 g for testosterone chicks; 58.66±4.63 g for control chicks, χ^2^=0.277, Holm-adjusted p=0.599). The effect of sex was not significant (marginal means±SE: males, 85.84±3.75 g; females, 83.45±3.59 g, F_1,15.32_=0.169, p=0.687), nor was the interaction effect of testosterone treatment and sex (F_1,16.40_=0.923, p=0.351).

**Figure 2.**
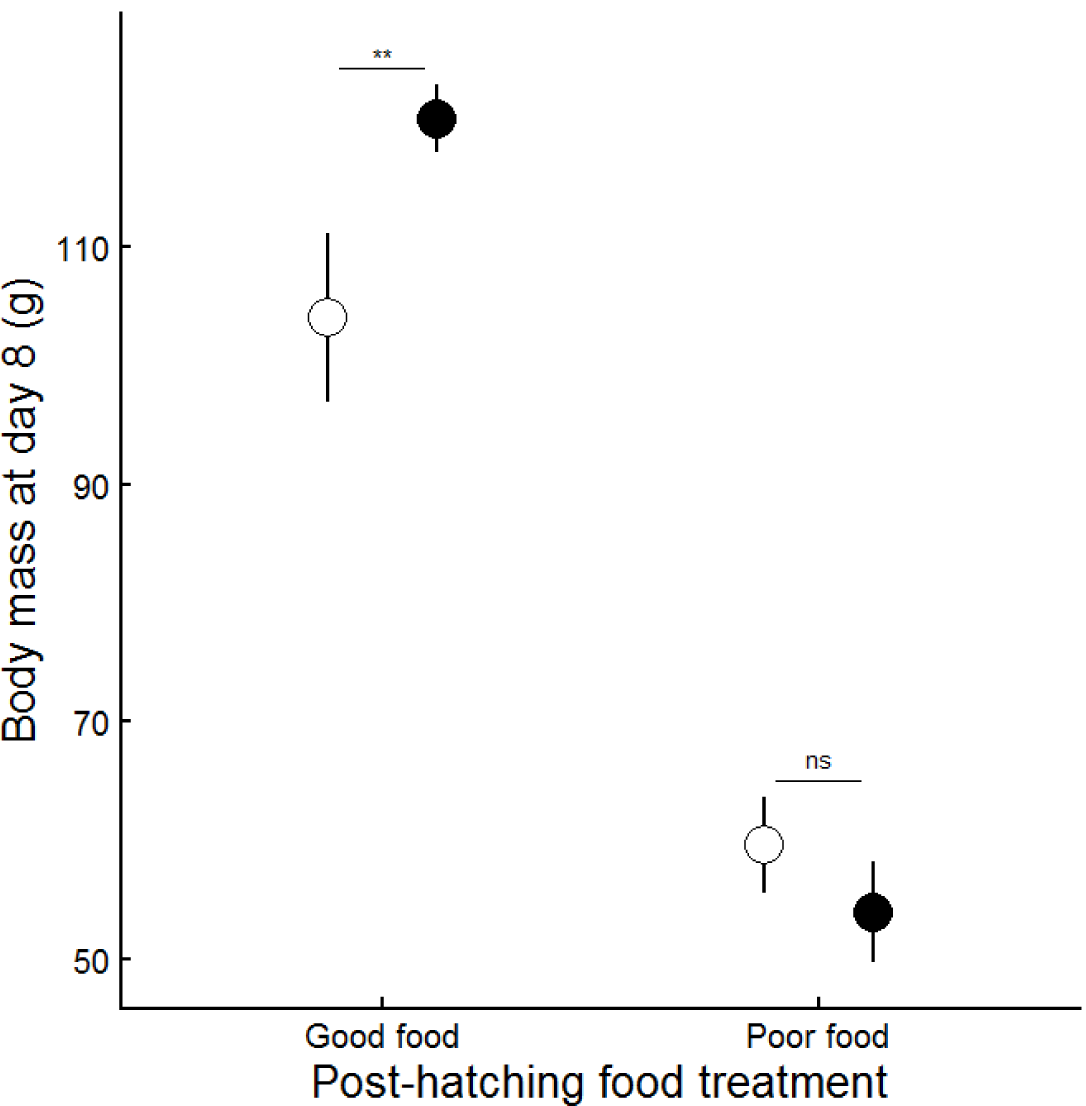
Means±SE of chick body mass at day 8 after hatching. Closed dots: chicks from testosterone-injected eggs; open dots: control chicks from vehicle-injected eggs. Post-hoc analysis: **, p < 0.01; ns, p > 0.05.

For body mass at day 26, when chicks were about to fledge, we found that of the chicks reared in the good food conditions, testosterone chicks were still on average 10.9 g (SE=5.60) heavier than control chicks, but the effect was not statistically significant (F_1,8.55_=3.788, p=0.085, Fig. 3). Neither sex nor egg mass showed significant effects at this age (p=0.312 and 0.397, respectively) and there was no interaction between testosterone treatment and sex (p=0.276).

**Figure 3.**
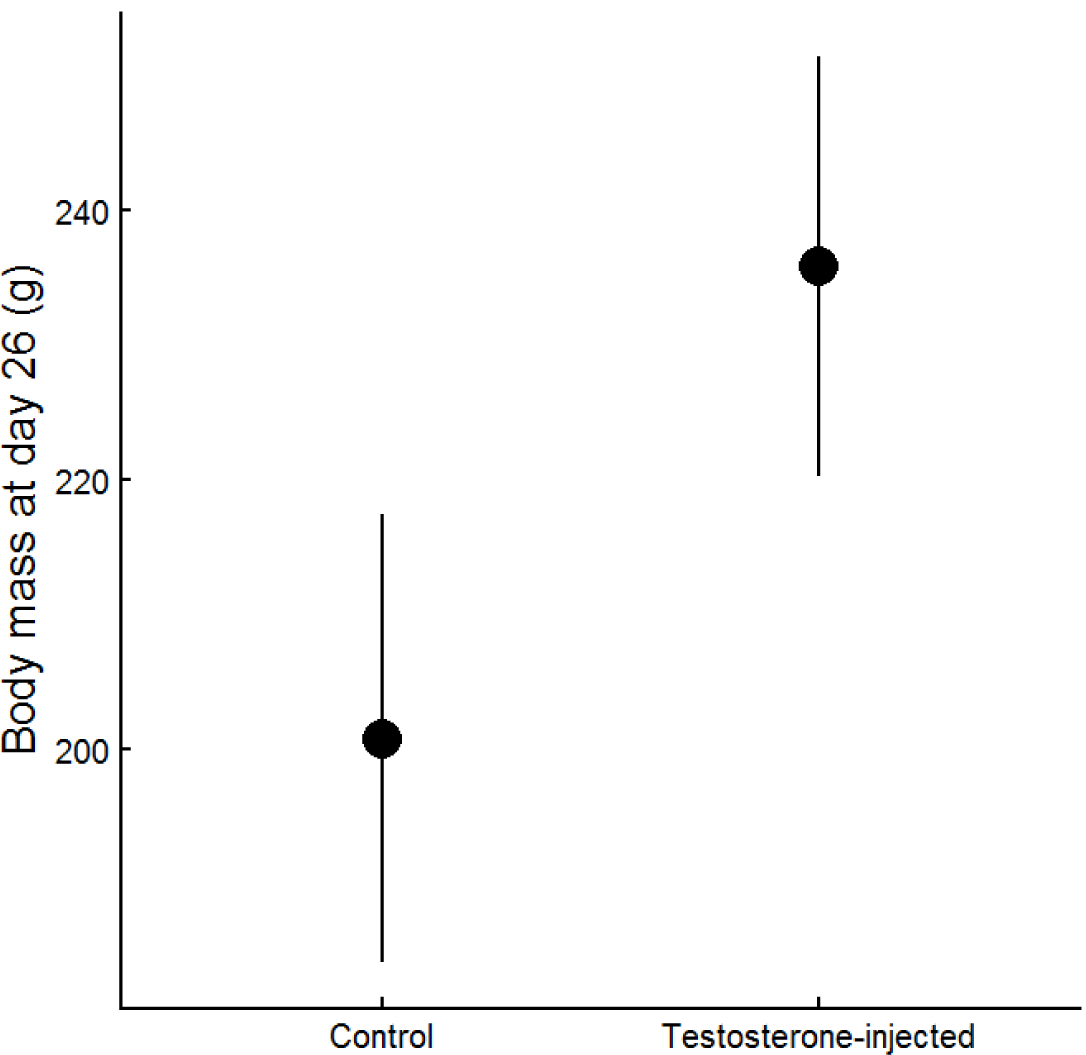
Mean ± SE of chick body mass at day 26, around fledging.

### Immunocompetence

Among the same-sex broods that survived to complete the SRBC tests (n=28), the testosterone fledglings showed significantly lower immune response against SRBC (n=13, marginal mean±SE = 2.05±0.75) than did control fledglings (n=15, marginal mean±SE = 4.08±0.68, t=-2.030, p=0.041, Fig. 4). Food restriction also showed a significant negative effect on the immune response (marginal means±SE = 5.18±0.56 for fledglings reared under the good food conditions, 0.95±0.91 for fledglings reared under the poor food conditions, t=-4.230, p<0.001, Fig. 4). The interaction between hormone and food treatment, however, did not have significant effects (p=0.396). There was no significant sex difference (p=0.101) or interaction between hormone treatment and sex (p=0.333).

**Figure 4.**
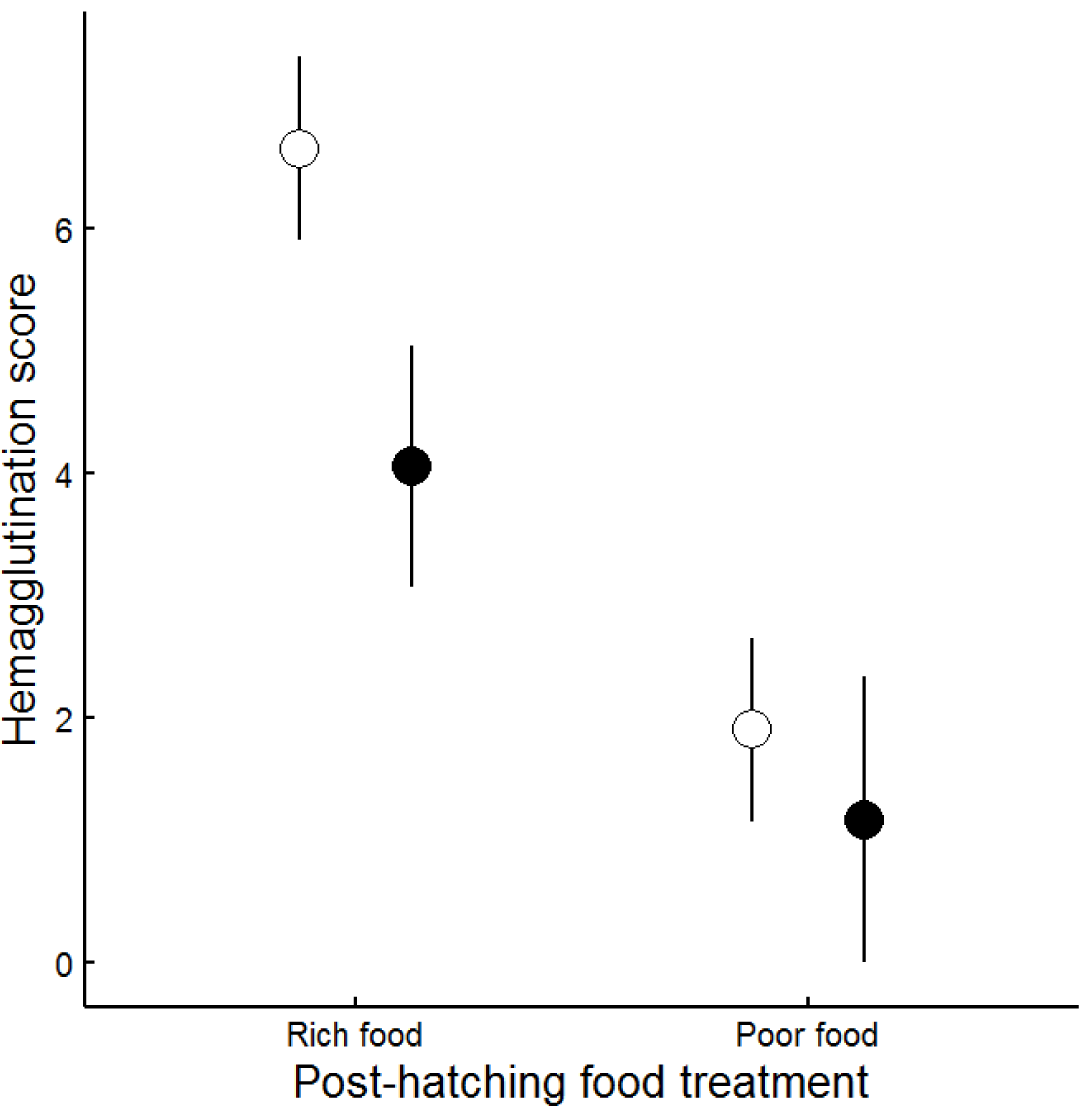
Mean ± SE of the score of anti-SRBC antibody titres from hemagglutination assay. Closed dots: chicks from testosterone-injected eggs; open dots: control chicks from vehicle-injected eggs.

## Discussion

Despite the fact that hatching asynchrony has been regarded as an adaptive maternal effect for optimizing the trade-off between offspring quantity and quality in different food conditions, maternal androgens in birds have been proposed as a tool for mothers to counteract the effects of hatching asynchrony^18,19^. This conceptual paradox, although put forward more than a decade ago^19^, still lacks sufficient effort to find a solution. The paradox could be resolved if maternal androgens have context-dependent effects, much like the context-dependent effects that hatching asynchrony has on survival of late-hatching offspring in good and poor food conditions. In this scenario, both maternal effects work together to downsize the brood when food is scarce by selectively hastening the death of the youngest offspring, and, when food is abundant, help rather than harm survival of late-hatching offspring by giving them, via maternal androgens, a boost in competing for food. Our findings support this idea: exposure to elevated yolk testosterone benefitted nestlings under good food conditions by enhancing growth (Fig. 2, 3), but caused higher mortality when food conditions were poor (Fig. 1). This is the first experimental evidence from a full factorial design that supports the notion of adaptive context-dependent effects of maternal hormones. It is also the first evidence that solves a long standing paradox in the field of hormone-mediated maternal effects, reconciling the seemingly opposing effects of hatching asynchrony and differential testosterone allocation to offspring. It shows that maternally engineered competitive asymmetries within broods arise from variation in yolk androgens and hatching asynchrony together, and allow parents to maximize the number of offspring they rear^26^. Moreover, in a system in which two maternal effects work in concert, mothers do not need to adjust the within-clutch pattern of testosterone deposition in relation to food abundance. As the effects of testosterone are context dependent, they can simply maintain higher testosterone concentrations in late-laid eggs regardless of pre egg-laying food conditions. This may explain why most studies did not find evidence for maternal adjustment of testosterone deposition in eggs in relation to food conditions, including in the pigeon^42^. Moreover, this system of competing asymmetries does not require parents to make assumptions about the future food conditions at the time of egg laying, long before the chicks hatch (in the pigeon 18 days).

The context dependent effects of yolk testosterone reported here may also explain earlier reported inconsistencies in the effects of *in ovo* injections of androgens^23^. Interestingly, the most cited study for the detrimental effects of yolk testosterone on survival, the study on American kestrels (*Falco sparverius*)^51^, was conducted in a poor food year (Sockman, personal communication). In contrast, a replication of that study on the closely related Eurasian kestrels (*F. tinnunculus*) in a good year found a positive effect (C. Dijkstra, J. Boonekamp and T.G.G. Groothuis, unpublished data). Similarly, a recently published field study on spotless starlings (*Sturnus unicolor*) reported that only among the second broods, when the food availability is generally deteriorated compared to when the parents raised their first broods, nestling mortality was significantly higher for the nestlings from androgen-treated eggs^52^. Another study by Cucco et al. (2009)^53^, who injected different doses of testosterone into grey partridge (*Perdix perdix*) egg yolks and supplemented the diet with β-carotene in half of the chicks from each testosterone treatment, found that supplemented β-carotene can remedy the immunosuppressive effects induced by higher levels of yolk testosterone. It is therefore important that future studies testing the effect of yolk hormones take into account the food conditions.

Our understanding of the mechanism underlying the context dependent effects of maternal yolk testosterone, however, remains to have several unanswered questions. The enhancing effects of testosterone on growth in good food conditions may have been due to an increase in begging behaviour, as has been found in several studies^54-56^, which will result in higher food delivery by the parents. Furthermore, the food dependent effects of elevated maternal testosterone on growth may also have been mediated by increases in basal metabolic rate (BMR), as elevated prenatal testosterone exposure increases BMR^27-29^. Indeed, during food supplementation, increased BMR has been shown to enhance growth, whereas during food restriction, increased BMR can depress growth^57,58^. Therefore, the higher mortality of chicks from testosterone-injected eggs observed in the poor food condition may have been the result of a higher, but unsatisfied, energy demand. Our findings showed that most nestling mortality occurred after 8 days of age (Fig. 1). In this species, the energy and nutrient demands of chicks most likely peak around 10 days after hatching, and the higher energy demand around that time may have induced a fatally negative energy balance of testosterone-injected chicks in the poor food conditions. Moreover, if prenatal testosterone increased begging behaviour in the poor food conditions, this would have further exacerbated energy loss and the parents would not have been able to compensate it.

Our results indicate that regardless of food conditions, testosterone-injected fledglings showed a weaker response to an immune challenge (injected sheep red blood cells). This effect of immunosuppression is consistent with previous studies in other avian species^30-34^. Although the interaction effect on chicks’ immune response between the yolk testosterone injection and food conditions was not significant, Figure 4 obviously suggests that a clear difference was only observed among chicks reared under good food conditions. This is likely due to a floor effect as food restriction induced already a very strong suppression of the immune response, and the suppressed responses would be difficult to differentiate as the antibody titers cannot go below 0. This also suggests that high resource inputs are required to maintain good immune function^59^. It is possible that the high rate of energy expenditure caused by maternal testosterone exposure, combined with the low rates of energy inputs due to poor food conditions, left insufficient resources for maintaining the immune system.

In conclusion, our results suggest the co-evolution of two maternal effects to optimize the final brood size that are adaptive for the mother but not necessarily optimal for all offspring. The context dependent effects of maternally deposited testosterone on chick survival can be used as an explanation for the apparent discrepancies in the literature on this subject that currently hamper progress in our understanding to the functions of maternal hormones. When these are taken into consideration, the field of hormone-mediated maternal effects may move forward substantially.

## Acknowledgement

We dedicate this paper to the memory of Dr. Cor Dijkstra, a beloved teacher, mentor, and friend, who peacefully left us in November, 2017. We greatly appreciate all animal care takers for their help, Roelie Veenstra-Wiegman for the help on sexing, Bernd Riedstra for the help on SRBC test, and Charlotte Deerenberg for valuable comments on the manuscript.

## Notes

### Competing Interest Statement

The authors have declared no competing interest.

