## Supplementary Material for "Co-evolved maternal effects selectively eliminate offspring depending on resource availability"

**Supplementary methods**

*Housing*

The pigeons used in this study came from our rock pigeon colony housed in a large outdoor aviary (45m long × 9.6 m wide × 3.75 m high). In early April 2012, they were re-housed in several smaller identical aviaries (4.01 m long × 1.67 m wide × 2.2 m high), with two pairs in each aviary, in the animal facility of the Centre of Life Sciences of the University of Groningen. Because their pair bonds can last for a lifetime (Johnston and Janiga 1995), we selected already established pairs when known, or randomly paired males with females if their partner identity was unknown. Established and random breeding pairs were assigned to the two food condition treatment groups in equal proportion. After re-housing them in the smaller aviaries, pigeons were given at least one week to acclimate before the experiment started. To initiate breeding, two nest boxes (60 cm long × 50 cm deep × 36 cm high) with nest materials and a nest bowl were placed into each small aviary. Identical nest-boxes were also provided in the large aviary to induce breeding in other pigeons, whose eggs were also collected and used for the experiment.

*Egg injections*

The testosterone solution for injection was prepared by dissolving crystalline testosterone (Fluka) in sterilized sesame oil at a concentration of 920 ng testosterone/ml oil, so that injecting 50 µl of this solution elevated the amount of testosterone in the yolks of the $1^{st}$-laid eggs (mean testosterone = 23 ng per yolk, n = 11) to the amount present in the $2^{nd}$-laid eggs (mean testosterone = 69 ng per yolk, n = 11), as measured by radioimmunoassay using a commercial kit ('Spectria 68628', Espoo, Finland, cross reactivity to A4 and 5α-DHT was 1.7% and 2.6 respectively, all others < 0.31%; recoveries averaged 82.2%; intra-assay CV=1.4%). We injected 50 µl testosterone solution or pure sesame oil into the egg yolk using a 0.5 ml U-100 insulin syringe [0.33 mm (29G) needle × 12.7 mm syringe, BD Micro-Fine™]. Eggs were first placed sideways for a few minutes, allowing the yolk to float up. Then the eggshell was punctured ~30° to 45° above from the blunt end by a needle attached to an insulin syringe filled with testosterone solution or sesame oil. After withdrawing the syringe, the hole in the eggshell was sealed by a small piece (~25mm$^2$) of artificial skin (Hansaplast ™).

*Chick measurements*

Body mass was measured using a digital scale with an accuracy of 1 g. Head-bill length and tarsus length were measured with digital callipers to the nearest 0.01 mm. Head-bill length was measured from the back of the skull to the tip of the bill. The length of the tarsometatarsal bone of the right leg was measured as ‘tarsus length’. Wing length was measured from the carpus to the tip of the skin (when chicks did not have primaries) of the right wing with a digital calliper to the closest 0.01 mm or to the tip of the longest primary feathers with a ruler to the closest 0.5 mm.

*SRBC test and hemagglutination assay*

We used sheep red blood cells (SRBC) as antigens to test the humoral immune response of pigeon fledglings. All SRBC used in this study were from one identical sheep and stored in Alsever’s solution (S.B.-0011, Harlan ™, UK). New SRBC in Alsever’s solution was washed with phosphate buffered saline (PBS) and diluted to 2% SRBC solution in PBS ($5\times10^8$ cells/ml). When chicks were 37-45 days old, every pigeon fledgling was immunized with 0.5 ml 2% SRBC solution via intraperitoneal injection. Two days prior to injection, pre-treatment blood samples (100-200 µl) were taken from the brachial vein. Six days after SRBC injection, at the day of peak antibody production, (Ros et al. 1997, Casagrande et al 2012), post-treatment blood samples were taken. Both blood samples were centrifuged to separate the plasma from blood cells. The plasma samples were then stored at -20°C until the hemagglutination assay.

The protocol of hemagglutination assay was based on Ros et al. (1997) to determine the titres of anti-SRBC antibodies. The frozen plasma samples were thawed under room temperature and placed in a water bath at 56°C for 30 min to remove proteins that will lyse the sheep red blood cells. Then 20 µl of the plasma was taken and diluted 1:1 in PBS in the first column of a U-shaped microtiter plates, and further diluted in a series of $2^2$, $2^3$, $2^4$… to $2^{12}$. Twenty µl of freshly-washed 2% SRBC solution was added to every well and all titre plates with samples were incubated at 37°C for one hour. Based on the presence of hemagglutination, the antibody titres were scored visually as the highest dilution of plasma by an experienced person who was blind to all experimental treatments. The scores were then represented as integers on a $\log_2$ scale. When the plasma from one sample was sufficient, we analysed a second duplicate and took the mean score from the two duplicates.

**Supplementary results**

**Results on chick head-bill, tarsus, and wing length**

*Day 8 after hatching:*

For head-bill length, the interaction between hormone injection and food conditions was approaching significance ($F_{1,15.61}$=3.553, p=0.078), showing a consistent trend with the interaction effect found in body mass (Fig. S2A). Sex difference was not significant ($F_{1,15.38}$=0.004, p=0.950), nor was the interaction between testosterone treatment and sex (p=0.107).

For tarsus length, the interaction between testosterone treatment and food conditions was also significant ($F_{1,15.84}$=4.911, p=0.042. Fig. S2B), consistent with the result in body mass. Post-hoc interaction analyses showed that in the good food condition, testosterone-chicks had significantly longer tarsi (marginal mean±SE = 27.59±0.47 mm) than control-chicks (marginal mean±SE = 26.41±0.47 mm, Holm-adjusted p = 0.035), while in the poor food condition there was no significant difference of tarsus length between testosterone- and control-chicks (marginal means±SE = 23.11±0.52 mm, 23.58±0.49 mm, respectively, Holm-adjusted p=0.394). Similarly, no sex difference ($F_{1,15.24}$=0.052, p=0.823) or sex-specific effect of testosterone treatment (p=0.521) was observed.

For wing length, the interaction between T treatment and food conditions was not significant ($F_{1,16.23}$=1.705, p=0.210), although it still showed similar pattern as in other biometric variables (Fig. S2C). Sex was also non-significant ($F_{1,15.10}$=0.303, p=0.590), nor was the interaction between T treatment and sex (p=0.759).

*Day 26 after hatching:*

At this age, much larger variations were observed in the three variables of skeletal growth. Testosterone-chicks still showed higher averages than control-chicks in tarsus and wing length (Fig. S3), but the difference was all non-significant (head-bill length, $F_{1,8.40}$=-0.043, p=0.840; tarsus length, $F_{1,8.53}$=1.437, p=0.263; wing length, $F_{1,8.72}$=2.628, p=0.141).

**Table S1.** Macronutrient components of grain mixtures in food treatment.

| Name of grain mixture | Tortelduivenvoer (KASPER™ 6721) | Sierduivenvoer (KASPER™ 6712) | Duivenkorrel[a] (KASPER™ P40) | Grain mixture with broken corns |
|---|---|---|---|---|
| Protein | 11.90% | 13.30% | 15.60% | 10.10% |
| Fat | 5.20% | 3.40% | 2.80% | 2.40% |
| Cellulose | 6% | 4.40% | 2.10% | 2.80% |
| Ash | 2.40% | 2.40% | 4% | 1.60% |

[a] also contains calcium 0.6%, potassium 0.6%, lysine 6g, sodium 0.1%, E 672 vitamin A 19000 lE/kg, vitamin E 100 lE/kg, E 671 vitamin D3 300 lE/kg, and copper 12mg/kg

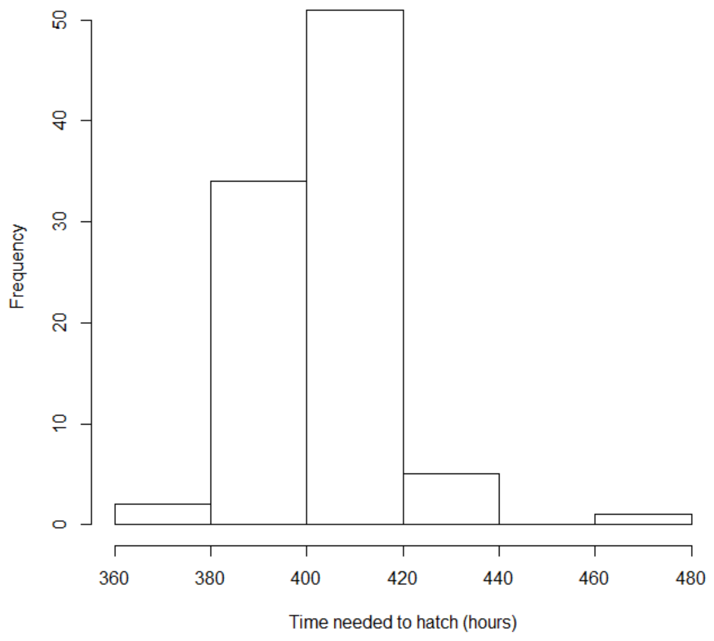


**Figure S1.** Histogram of the time from returning to hatch.

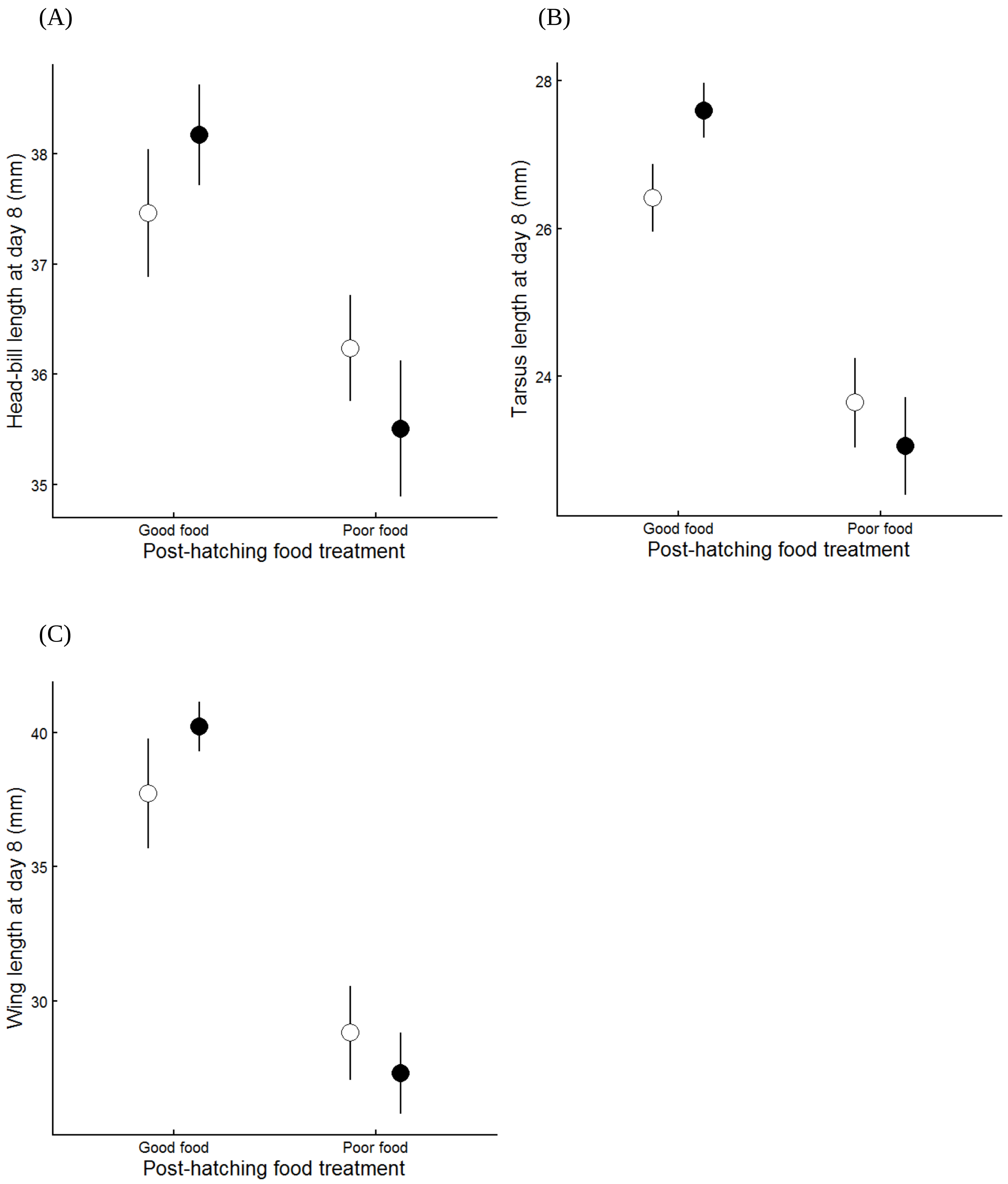


**Figure S2.** Means±SE of chick head-bill length (A), tarsus length (B), and wing length (C) at day 8 after hatching. Close dots: chicks from testosterone-injected eggs; open dots: chicks from vehicle-injected eggs.

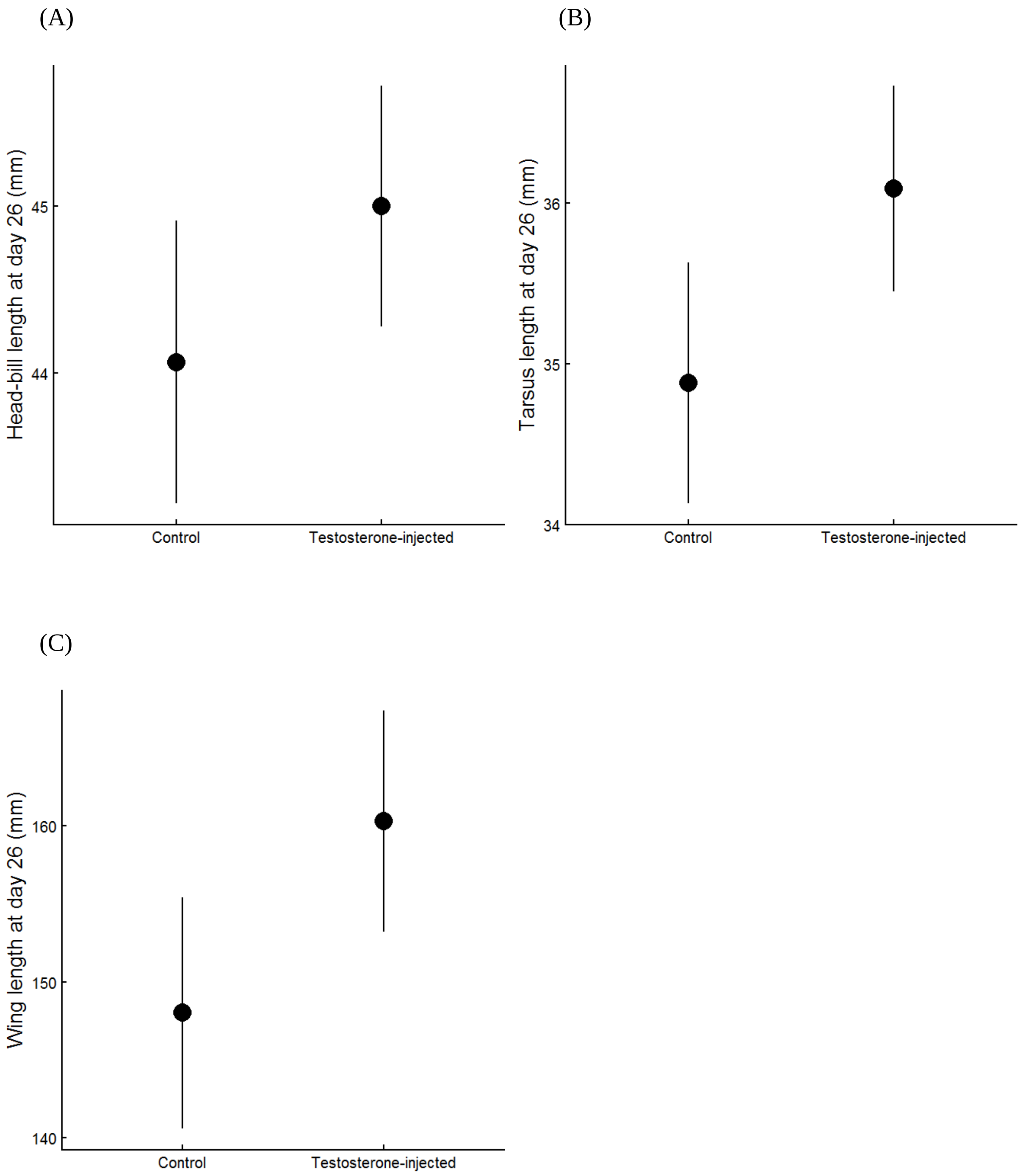


**Figure S3.** Mean±SE of chick head-bill length (A), tarsus length (B), and wing length (C) in the good food condition at day 26 after hatching.
